# The development of object recognition requires experience with the surface features of objects

**DOI:** 10.1101/2022.12.30.522302

**Authors:** Justin N. Wood, Samantha M. W. Wood

## Abstract

What role does visual experience play in the development of object recognition? Prior controlled-rearing studies suggest that newborn animals require slow and smooth visual experiences to develop object recognition. Here we examined whether the development of object recognition also requires experience with the surface features of objects. We raised newborn chicks in automated controlled-rearing chambers that contained a single virtual object, then tested their ability to recognize that object from familiar and novel viewpoints. When chicks were reared with an object that had surface features, the chicks developed view-invariant object recognition. In contrast, when chicks were reared with a line drawing of an object, the chicks failed to develop object recognition. The chicks reared with line drawings performed at chance level, despite acquiring over 100 hours of visual experience with the object. These results indicate that the development of object recognition requires experience with the surface features of objects.

## I. INTRODUCTION

Mature animals have powerful object recognition abilities. For example, after just a brief glimpse of an object, humans can recognize objects across substantial variation in the retinal images produced by the object, due to changes in viewpoint, size, illumination, and so forth (reviewed by DiCarlo, Zoccolan, & Rust, 2012). However, the origins of object recognition are still not well understood. What role does early visual experience play in the development of object recognition? Does the development of object recognition require a specific type of visual experience with objects?

Human infants are not well suited for addressing these questions because they cannot be raised in strictly controlled environments from birth. In contrast, controlled-rearing studies of newborn animals can directly probe the role of experience in development. By systematically manipulating the visual experiences provided to newborn animals and measuring the effects of those manipulations on behavioral and neural development, controlled-rearing studies can isolate the specific experiences that drive the development of object recognition.

Prior controlled-rearing studies with newborn chicks have revealed two types of experiences that are necessary for the development of object perception: slow and smooth experiences with objects (Prasad, Wood, & Wood, 2019; Wood, 2016; Wood & Wood, 2016; 2018; Wood, Prasad, Goldman, & Wood, 2016). When newborn chicks were reared with virtual objects that changed slowly and smoothly over time (akin to natural objects), the chicks successfully developed object recognition, including the ability to recognize objects across novel viewpoints, backgrounds, and motion speeds. Conversely, when chicks were reared with objects that moved too quickly or non-smoothly, the chicks failed to develop object recognition. Thus, without slow and smooth visual experiences, newborn chicks develop inaccurate object representations. Here, we extend these findings by examining whether the development of object recognition also requires experience with the surface features of objects. The term “surface features” refers to the features (e.g., color, texture, and shading) of the surfaces between the boundaries of an object.

There are mixed perspectives on the importance of surface features in object recognition. On one hand, a large number of studies have shown that human adults can readily recognize objects depicted in line drawings, which lack surface features such as color, texture, and shading (e.g., Biederman, 1987; Biederman & Ju, 1988; Ishai, Ungerleider, Martin, & Haxby, 2000; Walther, Chai, Caddigan, Beck, & Fei-Fei, 2011). This ability to perceive and understand line drawings emerges early in development. For instance, infants begin showing enhanced attention to lines that depict corners and edges in the first year of life (Yonas & Arterberry, 1994),^1^ and young children use lines to define the boundaries of objects in their first attempts to depict the world (Goodnow, 1977). Humans have also used line drawings to capture scenes since prehistoric times (Clottes, 2000; Kennedy & Ross, 1975). Furthermore, many nonhuman animals can understand line drawings. Chimpanzees can recognize objects presented in line drawings (Itakura, 1994; Tanaka, 2006) and pigeons can recognize line drawings of objects that are rotated in depth, even after exposure to just a single depth orientation (Wasserman et al., 1996). Even insects appear to use line representation to some extent in biomimicry (Sayim & Cavanagh, 2011). Together, these studies indicate that the ability to understand line drawings emerges early in development and is shared with a wide range of animals.

On the other hand, many studies provide evidence that surface features play an important role in object recognition (e.g., Hayward & Williams, 2000; Naor-Raz, Tarr, & Kersten, 2003; Rossion & Pourtois, 2004; Tarr, Kersten, & Bulthoff, 1998; Vuong, Peissig, Harrison, & Tarr, 2005; Wurm, Legge, Isenberg, & Luebker, 1993). Human adults can recognize an object faster when the light source remains in the same location compared to when the light source moves (Tarr et al., 1998), and surface features can affect the speed and accuracy of object recognition and object naming (Price & Humphreys, 1989). During their first months of life, human infants also rely on the motion of surface features to build object representations (reviewed by Spelke, 1990).

In all of the studies cited above, the subjects had acquired months to years of visual experience with real-world objects before they were tested. Thus, these studies do not reveal whether newborn brains can understand line drawings at the onset of vision or whether the development of this ability requires experience with natural visual objects. The present study distinguishes between these possibilities by testing whether newborn chicks can recognize objects presented in line drawings at the onset of vision, in the absence of prior visual experience with natural objects. Specifically, we contrasted the object recognition performance of newborn chicks reared with line drawings of objects versus realistic objects with surface features. The three experiments presented here allow for a direct test of the importance of surface features in the development of object recognition.

### A. Using automated controlled rearing to study the origins of object recognition

To examine the role of surface features in the development of object recognition, we used an automated controlled-rearing method (Wood, 2013). There are two benefits to using automated methods to probe the origins of visual intelligence. First, automation allows large amounts of precise behavioral data to be collected from each subject. In the present study, each chick’s behavior was recorded continuously (24/7) for up to two weeks, providing precise measurements of their object recognition performance. Second, since computers (rather than researchers) present the stimuli and code the behavior, automation eliminates the possibility of experimenter error and bias (Wood & Wood, 2019).

We used newborn chicks as an animal model because they are an ideal model system for studying the origins of object recognition (Wood & Wood, 2015). First, newborn chicks can be raised in strictly controlled environments immediately after hatching (e.g., environments containing no real-world objects). As a result, it is possible to control and manipulate all of the chicks’ visual object experiences from the onset of vision. Second, chicks imprint to objects seen in the first few days of life and will attempt to reunite with those objects when separated (Horn, 2004). This imprinting behavior emerges spontaneously and provides a reliable behavioral assay for measuring chicks’ object recognition abilities. Third, birds and mammals process sensory input using homologous cortical circuits with similar connectivity patterns (Güntürkün & Bugnyar, 2016; Jarvis et al., 2005; Karten, 2013; Sacho et al., 2020). Since birds and mammals use homologous neural circuits to perceive the world, controlled-rearing studies of newborn chicks can inform our understanding of the development of both avian and mammalian vision. Finally, newborn chicks develop high-level object recognition. For example, newborn chicks can solve the visual binding problem, building integrated object representations with bound color-shape features (Wood, 2014). Chicks can also parse objects from complex backgrounds (Wood & Wood, under review), build view-invariant object representations (Wood, 2013; 2015), and recognize objects rapidly, within a fraction of a second (Wood & Wood, 2017).

In Experiments 1-2, newborn chicks were reared for one week in strictly controlled environments that contained no objects other than a single virtual object (Figure 1A). For one group of chicks, the virtual object contained surface features (Surface Feature Condition), whereas for another group of chicks, the virtual object was a line drawing animation that lacked surface features (Line Drawing Condition). In the second week of life, we used a two-alternative forced-choice procedure to examine whether the chicks could recognize their imprinted object across familiar and novel viewpoints. If chicks can recognize objects presented in line drawings at the onset of vision, then their performance should be high in both conditions. Conversely, if the development of object recognition requires visual experience with the surface features of objects, then the chicks should develop more accurate object recognition abilities in the Surface Feature Condition than the Line Drawing Condition.

**Figure 1.**
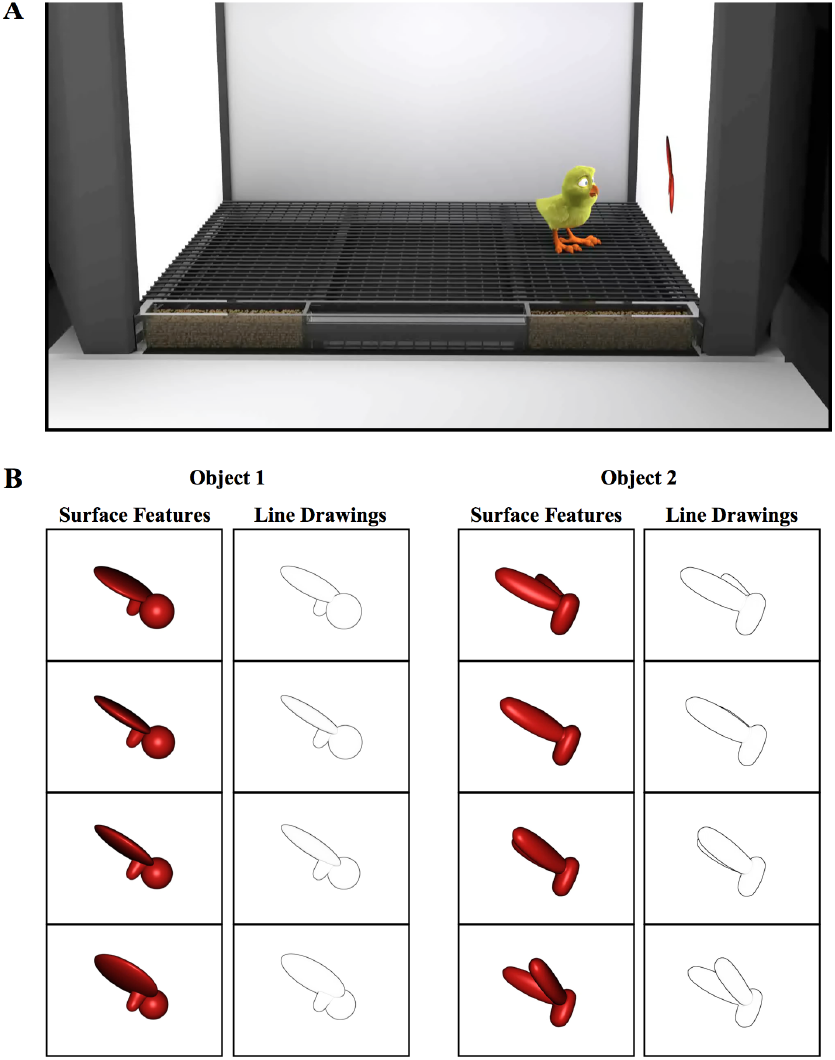
**(A)** Illustration of a controlled-rearing chamber. The chambers contained no real-world objects. To present object stimuli to the chicks, virtual objects were projected on two display walls situated on opposite sides of the chamber. During the input phase (1^st^ week of life), newborn chicks were exposed to a single virtual object either with surface features or without surface features (line drawing). The virtual objects.

## II. EXPERIMENT 1

In Experiment 1, newborn chicks were reared with a single virtual object moving through a limited 60° viewpoint range. In the test phase, we examined whether the chicks could recognize that object across 12 different viewpoint ranges. The chicks were either raised and tested with line drawings or with objects containing surface features. The text describing the methods is partly adapted from Wood (2013).^2^

### A. Method

#### Subjects

Twenty-three domestic chicks of unknown sex were tested. We tested 11 subjects in the Surface Feature Condition (Wood, 2013) and 12 subjects in the Line Drawing Condition. No subjects were excluded from the analyses. The eggs were obtained from a local distributor and incubated in darkness in an OVA-Easy incubator (Brinsea Products Inc., Titusville, FL). After hatching, we moved the chicks from the incubation room with the aid of night vision goggles. Each chick was placed, singly, in a controlled-rearing chamber. This research was approved by The University of Southern California Institutional Animal Care and Use Committee.

#### Controlled-Rearing Chambers

The controlled-rearing chambers (66 cm length × 42 cm width × 69 cm height) were constructed from white, high-density plastic. Each chamber contained no real-world (solid, bounded) objects (Figure 1A). We presented object stimuli to the chicks by projecting animations of virtual objects on two display walls situated on opposite sides of the chamber. The display walls were 19” liquid crystal display (LCD) monitors with 1440 × 900 pixel resolution. The monitors had a refresh rate of 30-80 Hz. We provided food and water in transparent troughs in the ground (66 cm length × 2.5 cm width × 2.7 cm height). We fed the chicks grain because grain does not behave like an object (i.e., a heap of grain does not maintain a solid, bounded shape). The floors were wire mesh and supported 2.7 cm off the ground by transparent beams.

We embedded micro-cameras in the ceilings of the chambers to record all of the chicks’ behavior (9 samples/second, 24 hours/day, 7 days/week). We used automated image-based tracking software (EthoVision XT, Noldus Information Technology, Leesburg, VA) to track their behavior throughout the experiment. This automated data collection approach allowed us to collect 168 trials from each chick. In total, 7,728 hours of video footage (14 days × 24 hours/day × 23 subjects) were collected for Experiment 1.

#### Procedure

In the first week of life (input phase), newborn chicks were reared in controlled-rearing chambers that contained a single virtual object. On average, the object measured 8 cm (length) × 7 cm (height) and was displayed on a uniform white background. Eleven of the chicks were imprinted to Object 1 (with Object 2 serving as the unfamiliar object), and 12 of the chicks were imprinted to Object 2 (with Object 1 serving as the unfamiliar object). The objects were modeled after those used in previous studies that tested for invariant object recognition in adult rats (Zoccolan, Oertelt, DiCarlo, & Cox, 2009).

The object moved continuously (24 frames/s), rotating through a 60° viewpoint range about a vertical axis passing through its centroid (Figure 1B). The object only moved along this 60° trajectory; the chicks never observed the object from any other viewpoint in the input phase. The object switched display walls every 2 hours (following a 1-minute period of darkness), appearing for an equal amount of time on the left and right display wall. In the Surface Feature Condition, the imprinted object had realistic surface features, whereas in the Line Drawing Condition, the imprinted object was a line drawing animation of the object (Figure 1B, see Movie S1 for animations).

In the second week of life (test phase), the chicks received 168 test trials (24 test trials per day). During the test trials, the imprinted object was shown on one screen and an unfamiliar object was shown on the other screen. We expected the chicks to spend a greater proportion of time in proximity to the object that they perceived to be their imprinted object. The image-based tracking software scored the chick as being in proximity to an object when the chick occupied a 22 × 42 cm zone next to the object.

For all of test trials, the unfamiliar object was presented from the same viewpoint range as the imprinted object shown during the input phase. The unfamiliar object had a similar size, color, motion speed, and motion trajectory as the imprinted object from the input phase. Consequently, for all of the novel viewpoint ranges, the unfamiliar object was more similar to the imprinting stimulus (from a pixel-wise perspective) than the imprinted object was to the imprinting stimulus (for details see Wood, 2013). To recognize their imprinted object, the chicks needed to generalize across large, novel, and complex changes in the object’s appearance on the retina.

The chicks were tested across 12 viewpoint ranges (11 novel, 1 familiar). Each viewpoint range was tested twice per day. The test trials lasted 20 minutes and were separated from one another by 40-minute rest periods. During the rest periods, the animation from the input phase appeared on one display wall and a white screen appeared on the other display wall. The 12 viewpoint ranges were tested 14 times each within randomized blocks over the course of the test phase. Figure 2A illustrates how the objects were presented across the display walls during the input phase and test phase. In the Surface Feature Condition, the chicks were tested with objects containing surface features, whereas in the Line Drawing Condition, the chicks were tested with line drawings of the objects. In both conditions, the test objects moved continuously through a 60° viewpoint range.

**Figure 2:**
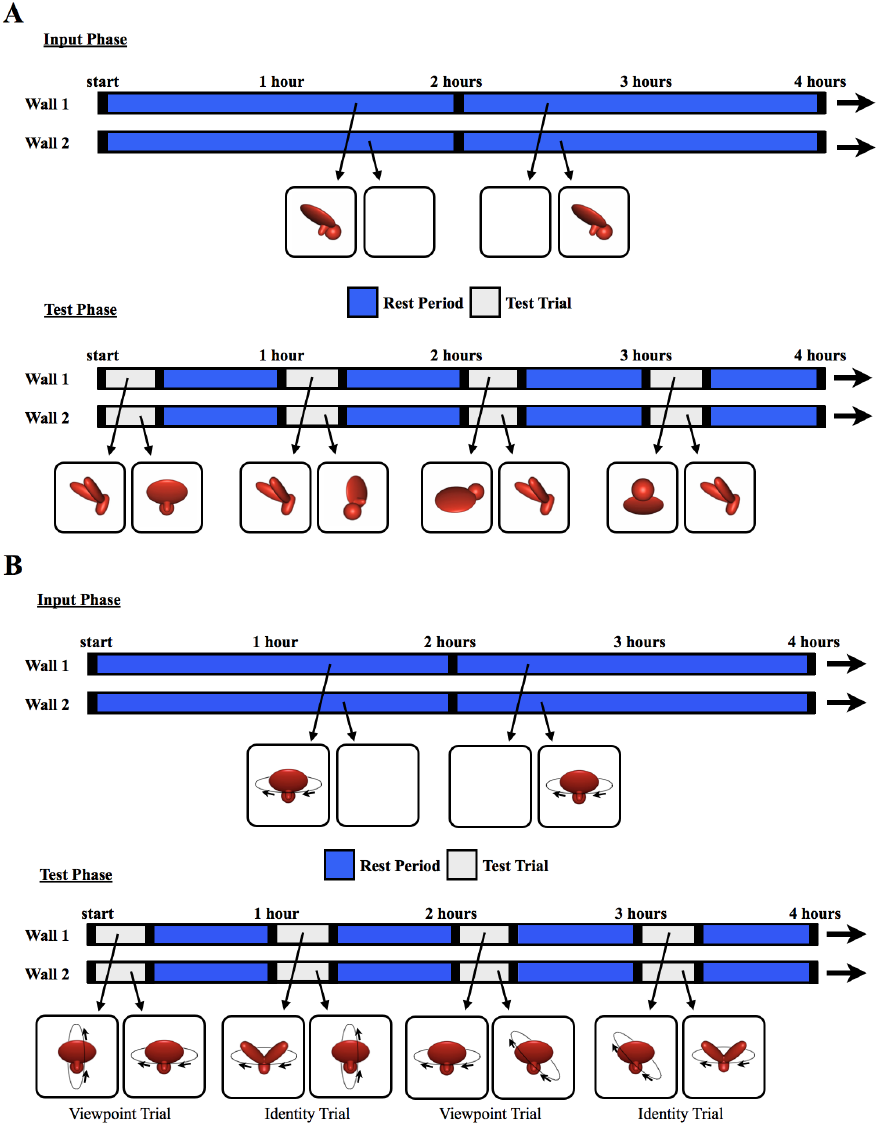
The experimental procedure. The schematics illustrate how the objects were presented for sample 4-hour periods during Experiment 1 and **(B)** Experiment 2. During the input phase, chicks were exposed to a single virtual object moving through a 60° (Experiment 1) or 360° (Experiment 2) viewpoint range. The object appeared on one wall at a time (indicated by blue segments on the timeline), switching walls every 2 hours, after a 1-min period of darkness (black segments). During the test trials, two virtual objects were shown simultaneously, one on each wall, for 20 minutes per hour (gray segments). The illustrations below the timeline are examples of paired test objects displayed in four of the test trials. Each test trial was followed by a 40-min rest period (blue segments). During the rest periods, the animation from the input phase was shown on one wall, and the other wall was blank. This figure shows the stimuli from the Surface Feature Condition. In the Line Drawing Condition, the chicks were raised and tested with line drawings rather than objects with surface features.

### B. Results

The results are depicted in Figure 3. For each viewpoint range, we computed the percent of time the chick spent with the imprinted object versus the unfamiliar object. Recognition performance exceeded chance level in the Surface Feature Condition (*t*(10) = 9.75, *p* < 10^−5^, Cohen’s *d* = 2.94), but did not exceed chance levels in the Line Drawing Condition (*t*(11) = 1.53, *p* = .15, Cohen’s *d* = .44). A repeated measures ANOVA with Viewpoint Range as a within-subjects factor and Condition (Surface Feature vs. Line Drawing) as a between-subjects factor revealed a significant main effect of Viewpoint Range (*F*(6.94, 145.81) = 2.73, *p* = .01, η _p_^2^ = .12) and Condition (*F*(1,21) = 52.13, *p* < .001, η_p_ ^2^ = .71). The interaction was also significant (*F*(6.94, 145.81) = 2.70, *p* = .01, η _p_^2^= .11). Recognition performance was significantly higher in the Surface Feature Condition than the Line Drawing Condition, both in terms of overall recognition performance (*t*(21) = 7.22, *p* < 10^−6^, Cohen’s *d* = 2.99; Figure 3A) and for each of the 12 viewpoint ranges (all *p*s < .05, Figure 3B).

**Figure 3:**
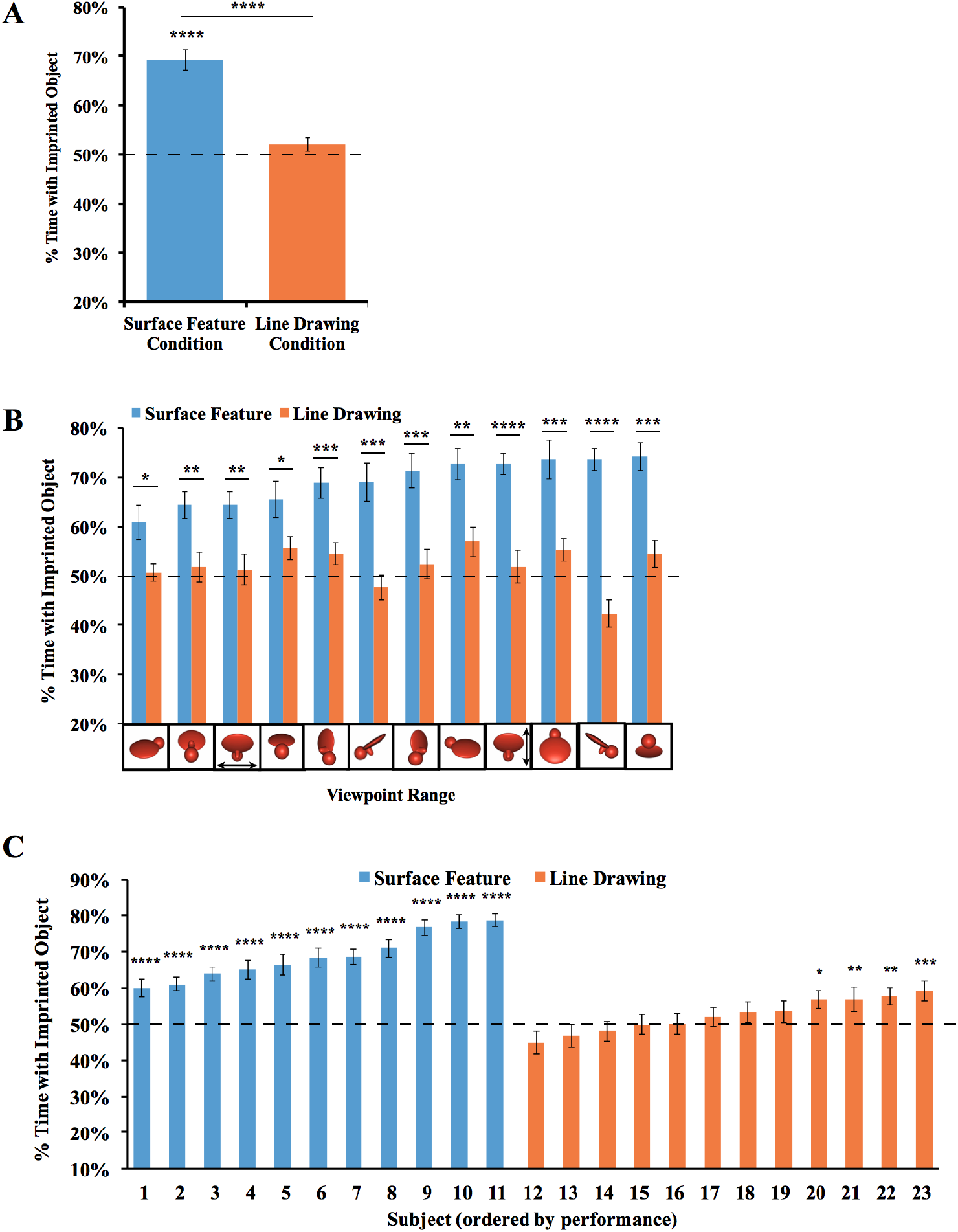
Results from Experiment 1. **(A)** Overall object recognition performance across the test phase. **(B)** Recognition performance on each of the 12 viewpoint ranges. **(C)** Recognition performance of each individual subject. The graphs show the percent of time spent with the imprinted object versus unfamiliar object. The dashed lines indicate chance performance. Error bars denote ±1 standard error. Asterisks denote statistical significance: \**P* < 0.05; \*\**P* < 0.01; \*\*\**P* < 0.001; \*\*\*\**P* < 0.0001 (two-tailed *t*-tests).

We also examined performance for each of the two imprinted objects. When the chicks were imprinted to Object 1 (see Figure 1 for reference), performance exceeded chance level in the Surface Feature Condition (*t*(4) = 6.54, *p* = .003, Cohen’s *d* =2.92), but not in the Line Drawing Condition (*t*(5) = 1.35, *p* = .24, Cohen’s *d* = -0.55). Performance was also significantly higher in the Surface Feature Condition than the Line Drawing Condition for Object 1 (*t*(9) = 7.11, *p* = .00006, Cohen’s *d* = 4.13). When the chicks were imprinted to Object 2, performance exceeded chance level in both the Surface Feature Condition (*t*(5) = 7.75, *p* = .001, Cohen’s *d* = 3.16) and the Line Drawing Condition (*t*(5) = 5.23, *p* = .003, Cohen’s *d* = 2.13), although performance was significantly higher in the Surface Feature Condition than the Line Drawing Condition (*t*(10) = 4.55, *p* = .001, Cohen’s *d* = 2.63). In general, newborn chicks developed superior object recognition abilities when reared with objects containing surface features versus line drawings.

Since over 100 test trials were collected from each chick, we could also measure each chick’s object recognition performance with high precision. As shown in Figure 3C, all chicks in the Surface Feature Condition successfully created view-invariant object representations (all *P*s < .0001). Conversely, only four of the 12 chicks in the Line Drawing Condition performed above chance level in the task, and those four subjects performed much worse than the subjects in the Surface Feature Condition.

### C. Discussion

In Experiment 1, newborn chicks developed enhanced object recognition performance when reared with objects containing surface features versus line drawings. Overall, the chicks reared with the line drawings performed at chance level, despite acquiring over 100 hours of visual experience with the line drawings during the input phase. Thus, the development of object recognition in newborn chicks requires visual experience with the surface features of objects.

To verify this conclusion under different testing conditions, we performed a second experiment with two key changes. First, rather than presenting the object from a 60° viewpoint range, the object moved through a 360° viewpoint range. As a result, the chicks were exposed to six times as many unique views of the object during the input phase. Second, we measured each chick’s object recognition abilities with Identity Trials and Viewpoint Trials (Figure 2B). The Identity Trials tested whether the chicks built object representations that were selective for object identity and tolerant to changes in viewpoint. The Viewpoint Trials tested whether the chicks built object representations that were selective for familiar viewpoints. The Identity Trials therefore tested the chicks’ view-invariant object recognition abilities, whereas the Viewpoint Trials tested whether the chicks could use an image-based matching strategy to recognize their imprinted object.

## III. EXPERIMENT 2

### A. Method

The text describing the methods is partly adapted from Wood & Wood (2016). The methods were identical to those used in Experiment 1, except in the following ways. First, 20 different subjects were tested. Ten chicks were tested in the Surface Feature Condition (Wood & Wood, 2016) and 10 chicks were tested in the Line Drawing Condition. No subjects were excluded from the analyses. Second, the imprinted object completed a 360° rotation every 15s around a frontoparallel vertical axis (see SI Movie 2 for animations). Third, the chicks were tested with Viewpoint Trials and Identity Trials. On the Viewpoint Trials, one display wall showed familiar viewpoints of the imprinted object (rotation around the familiar axis), whereas the other display wall showed novel viewpoints of the imprinted object (rotation around a novel axis, Figure 2B). If the chicks created object representations that were selective for familiar viewpoints, then they should have preferred the imprinted object rotating around the familiar axis over the novel axis. On the Identity Trials, one display wall showed the imprinted object rotating around a novel axis, whereas the other display wall showed a novel object rotating around the familiar axis (Figure 2B). Thus, to recognize their imprinted object on Identity Trials, the chicks needed to build view-invariant representations that were selective for object identity and tolerant to viewpoint changes.

The chicks received 24 test trials per day (168 test trials in total). Figure 2B shows how the objects were presented on the display walls during the input phase and test phase. In total, 6,720 hours of video footage (14 days × 24 hours/day × 20 subjects) were collected for Experiment 2.

### B. Results and Discussion

The results are shown in Figure 4. An ANOVA with the within-subjects factor of Trial Type (Viewpoint Trials vs. Identity Trials) and the between-subjects factor of Condition (Surface Feature vs. Line Drawing) revealed a significant main effect of Condition (*F*(1,18) = 48.55, *p* < .001, η_p_ ^2^ = .73), reflecting higher performance in the Surface Feature condition. The ANOVA also showed a significant main effect of Trial Type (*F*(1,18) = 14.01, *p* = .001, η_p_ ^2^ = .44), reflecting higher performance on the Identity Trials. The interaction was not significant (*F*(1,18) = 1.22, *p* = .29, η _p_^2^ = .06). In the Surface Feature Condition, performance was above chance level on the Identity Trials (one-sample *t*-test, *t*(9) = 7.84, *p* < .001, Cohen’s *d* = 2.48), but not on the Viewpoint Trials (*t*(9) = 1.41, *p*= .19, Cohen’s *d* = .45). In the Line Drawing Condition, performance did not exceed chance level on the Identity Trials (*t*(9) = 0.68, *p* = .51, Cohen’s *d* = .22) or the Viewpoint Trials (*t*(9) = 2.22, *p* = .053, Cohen’s *d* = .70; a “marginal” preference for novel viewpoints, the opposite direction of the expected response for object recognition). Thus, when chicks were reared with an object containing surface features, the chicks built object representations that were highly sensitive to identity features. When chicks were reared with line drawings, they did not show evidence for sensitivity to identity or viewpoint features.

**Figure 4:**
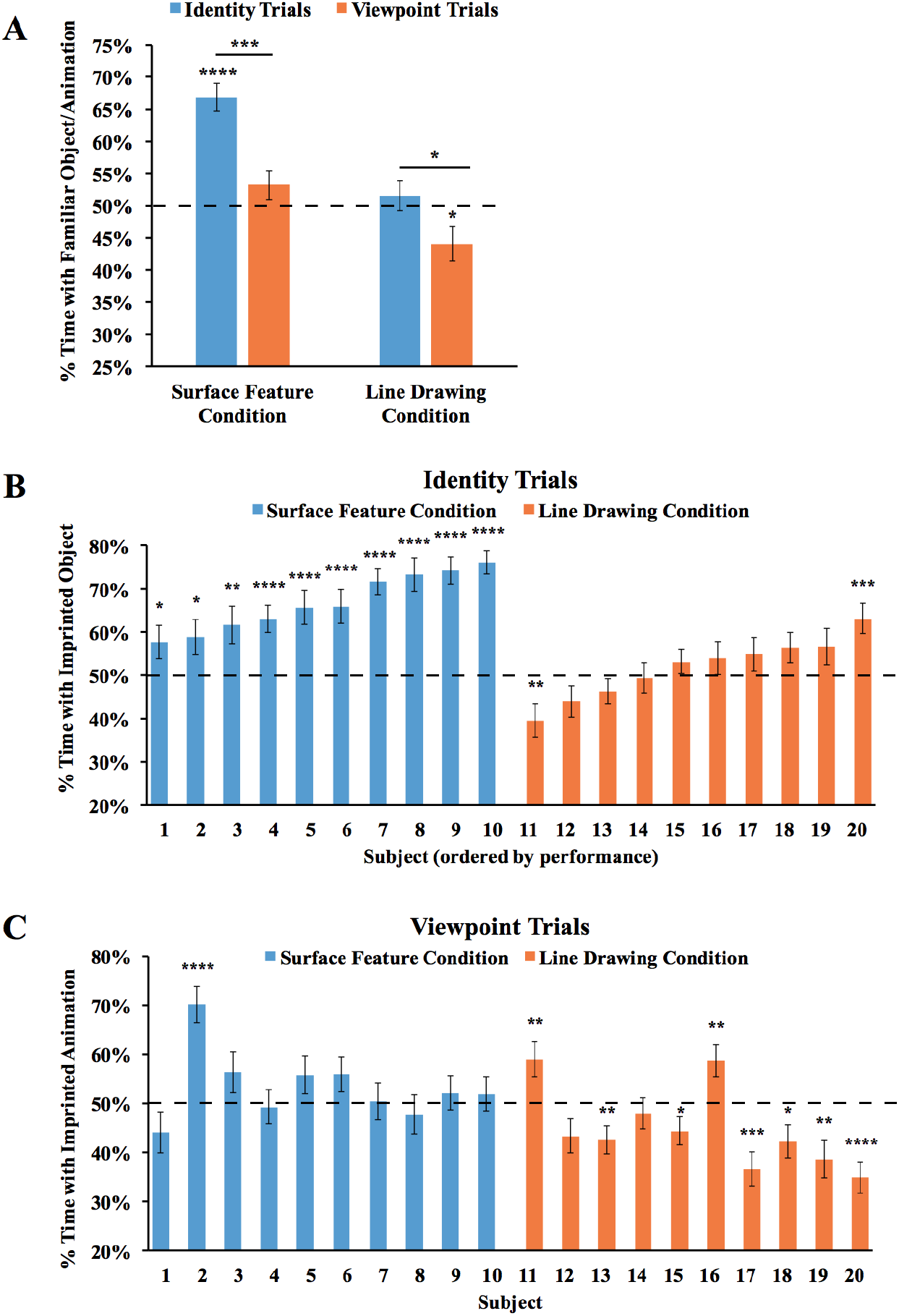
Results from Experiment 2. **(A)** Overall object recognition performance across the test phase. **(B)** Recognition performance of each individual subject on the Identity Trials. **(C)** Recognition performance of each individual subject on the Viewpoint Trials. The dashed lines indicate chance performance. Error bars denote ±1 standard error. Asterisks denote statistical significance: \**P* < 0.05; \*\**P* < 0.01; \*\*\**P* < 0.001; \*\*\*\**P* < 0.0001 (two-tailed *t*-tests).

We also examined performance for each of the two imprinted objects. We repeated the ANOVA above, but with the addition of Object as a main effect. The ANOVA revealed the same significant effects as before (significant main effects of Condition and Trial Type), and the main effect of Object was not significant, nor were the interactions (all *P*s > .3). When the chicks were imprinted to Object 1, performance exceeded chance level on the Identity Trials in the Surface Feature Condition (*t*(3) = 4.03, *p* = .03, Cohen’s *d* = 2.02), but not in the Line Drawing Condition *t*(5) = 0.16, *p* = .88, Cohen’s *d* = 0.06). On the Identity Trials, performance was significantly higher in the Surface Feature Condition than the Line Drawing Condition (*t*(8) =3.98, *p* = .004, Cohen’s *d* = 2.41). Similarly, when the chicks were imprinted to Object 2, performance exceeded chance level on the Identity Trials in the Surface Feature Condition (*t*(5) = 6.44, *p* = .001, Cohen’s *d* = 2.63), but not in the Line Drawing Condition (*t*(3) = 0.67, *p* = .55, Cohen’s *d* = 0.33). Again, on the Identity Trials, performance was significantly higher in the Surface Feature Condition than the Line Drawing Condition (*t*(8) =2.68, *p* = .03, Cohen’s *d* = 1.64). On the Viewpoint Trials, performance did not exceed chance level when the chicks were imprinted to Object 1 or Object 2 in the Surface Feature Condition or the Line Drawing Condition (*ps >* .15).

As shown in Figure 4B, all of the chicks in the Surface Feature Condition exceeded chance level on the Identity Trials (2 chicks, *P* < .05; 1 chick, *P* < .01; 7 chicks, *P* < .0001). In contrast, recognition performance was low for all of the chicks in the Line Drawing Condition. Only one chick exceeded chance level on the Identity Trials, while one other chick performed significantly below chance levels. As in Experiment 1, newborn chicks developed superior object recognition abilities when reared with objects containing surface features versus line drawings.

### C. Measuring the strength of the imprinting response in Experiments 1 & 2

To test the strength of the chicks’ imprinting response, we performed two additional analyses. First, we examined the proportion of time that the chicks spent in proximity to their imprinted object during the input phase. As shown in Figure 5A, the chicks in both Experiments 1 and 2 spent the majority of their time in proximity to the imprinted object during the input phase (Experiment 1 subjects imprinted to surface feature objects: *t*(10) = 40.47, *p* < 10^−11^, *d* = 12.20; Experiment 1 subjects imprinted to line drawings: *t*(11) = 6.27, *p* = .00006, *d* = 1.81; Experiment 2 subjects imprinted to surface feature objects: *t*(9) = 20.55, *p* < 10^−8^, *d* = 6.50; Experiment 2 subjects imprinted to line drawings: *t*(9) = 10.84, *p* = .000002, *d* = 3.43). Thus, the chicks successfully imprinted to both the line drawings and the objects with surface features. In both experiments, however, the imprinting response was stronger in the Surface Feature Condition than the Line Drawing Condition (Experiment 1: *t*(13.16) = 6.22, *p* = .00003, *d* = 2.55; Experiment 2: *t*(18) = 2.15, *p* = .05, *d* = 0.96), suggesting that the chicks imprinted less strongly to the line drawings than to the objects with surface features.

**Figure 5:**
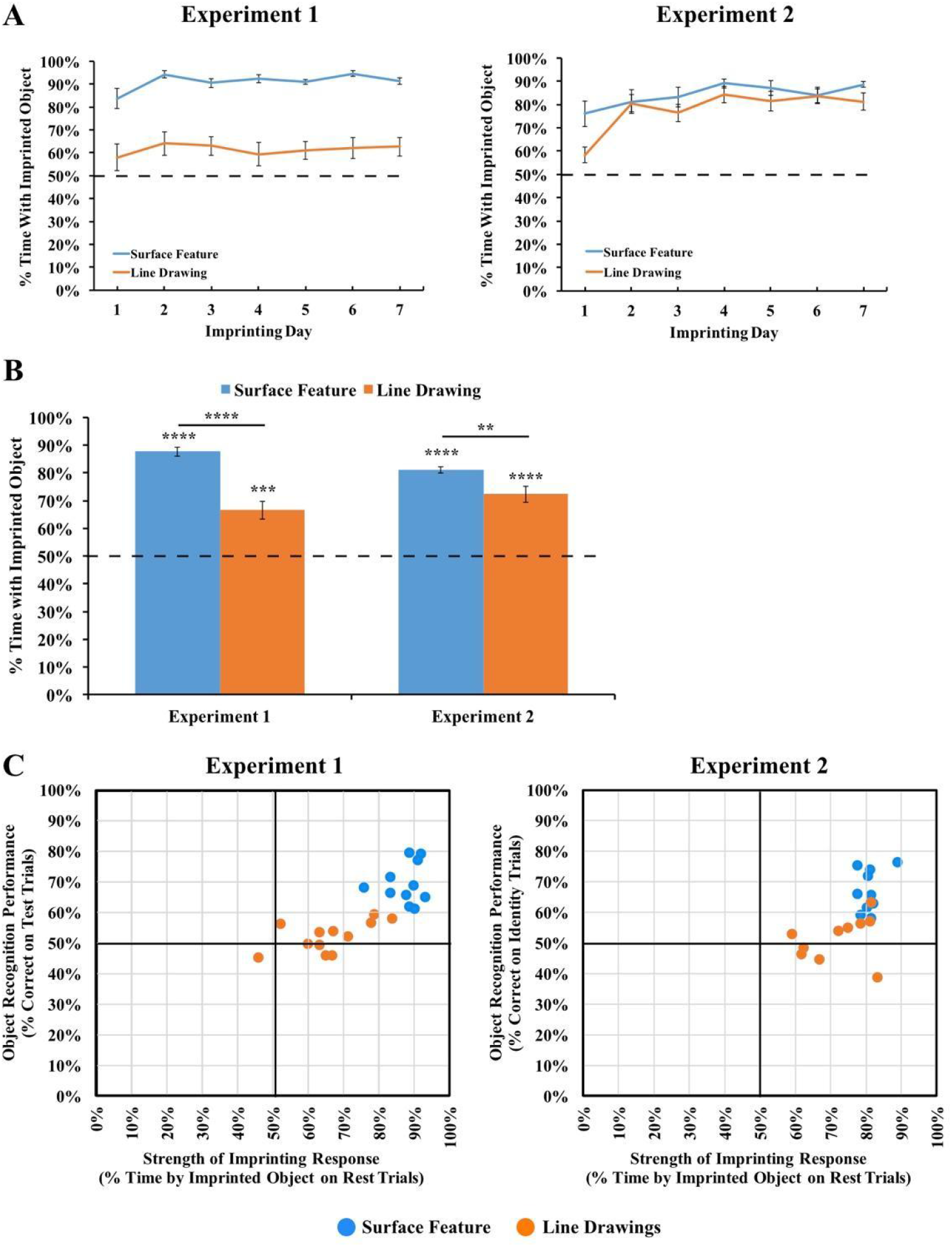
**(A)** Strength of the imprinting response in Experiments 1 and 2 during the input phase. **(B)** Strength of the imprinting response in Experiments 1 and 2 during the rest periods of the test phase. The dashed lines indicate chance performance. Error bars denote ±1 standard error. Asterisks denote statistical significance: \**P* < 0.05; \*\**P* < 0.01; \*\*\**P* < 0.001; \*\*\*\**P* < 0.0001 (two-tailed *t*-tests). The chicks successfully imprinted to both the line drawings and the objects with surface features, and this effect was stronger for the objects with surface features. **(C)** Comparison of the chicks’ object recognition performance and the strength of their imprinting response. The chicks developed enhanced object recognition performance when reared with objects with surface features compared to line drawings, even when the chicks imprinted to the objects at similar strengths.

Second, we examined the proportion of time the chicks spent with their imprinted object during the rest periods. During the rest periods, the imprinted object was presented on one display wall while the other display wall was blank. The rest periods therefore provided a measure of the strength of the chick’s attachment to the imprinted object during the test phase. As shown in Figure 5B, the chicks in both experiments spent the majority of their time in proximity to the imprinted object during the rest periods (Experiment 1 subjects imprinted to surface feature objects: *t*(10) = 24.91, *p* < 10^−9^, *d* = 7.51; Experiment 1 subjects imprinted to line drawings: *t*(11) = 5.35, *p* = .0002, *d* = 1.54; Experiment 2 subjects imprinted to surface feature objects: *t*(9) = 30.14, *p* < 10^−9^, *d* = 9.53; Experiment 2 subjects imprinted to line drawings: *t*(9) = 7.76, *p* = .00003, *d* = 2.45). However, the imprinting response was stronger in the Surface Feature Condition than the Line Drawing Condition (Experiment 1: *t*(21) = 5.91, *p* = .000007, Cohen’s *d* = 2.50; Experiment 2: *t*(11.30) = 2.89, *p* = .01, Cohen’s *d* = 1.29), providing additional evidence that the chicks imprinted less strongly to the line drawings than to the objects with surface features. Importantly, this reduction in the strength of the imprinting response cannot fully explain the low recognition performance because even the chicks that imprinted strongly to the line drawings still built inaccurate object representations (Figure 5C). Together, these analyses suggest that when chicks are reared with line drawings versus objects with surface features, the chicks develop an impaired imprinting response and build less accurate object representations.

## IV. EXPERIMENT 3

Experiments 1 and 2 indicate that the development of object recognition requires experience with the surface features of objects. However, there are three limitations to these results. First, the line drawings and the objects with surface features differed in several respects, including their color, contrast, hue homogeneity, and complexity. Thus, it is unclear which particular features caused the observed differences in recognition performance across the conditions. Indeed, there is extensive evidence that color is one of the most distinctive features encoded in imprinting (e.g. Bateson & Jaeckel, 1976; Ham & Osorio, 2007; Johnson et al. 1985; Miura et al., 2020; Nakamori et al. 2013; Wood, 2014), which raises the possibility that color differences may have influenced performance across the conditions. Second, Experiments 1 and 2 tested chicks’ recognition performance with the same two objects, so it is unclear whether these results generalize to other objects. Third, the imprinting response was less strong in the Line Drawing Condition than the Surface Feature Condition, potentially because of the color differences across objects. Given that recognition performance in this task is directly constrained by the strength of the imprinting response, the weaker imprinting response in the Line Drawing Condition likely produced lower recognition performance on the test trials.

To provide a more direct comparison of chicks’ recognition performance across conditions, we performed a third experiment in which the objects in the Surface Feature and Line Drawing Conditions were the same color (red). We also ensured that the objects did not have shadows and regions with different luminance values (as in Experiments 1 and 2), by using two-dimensional objects, rather than three-dimensional objects. The only difference between the conditions was whether the objects did, or did not, have surface features (Figure 6A).

**Figure 6:**
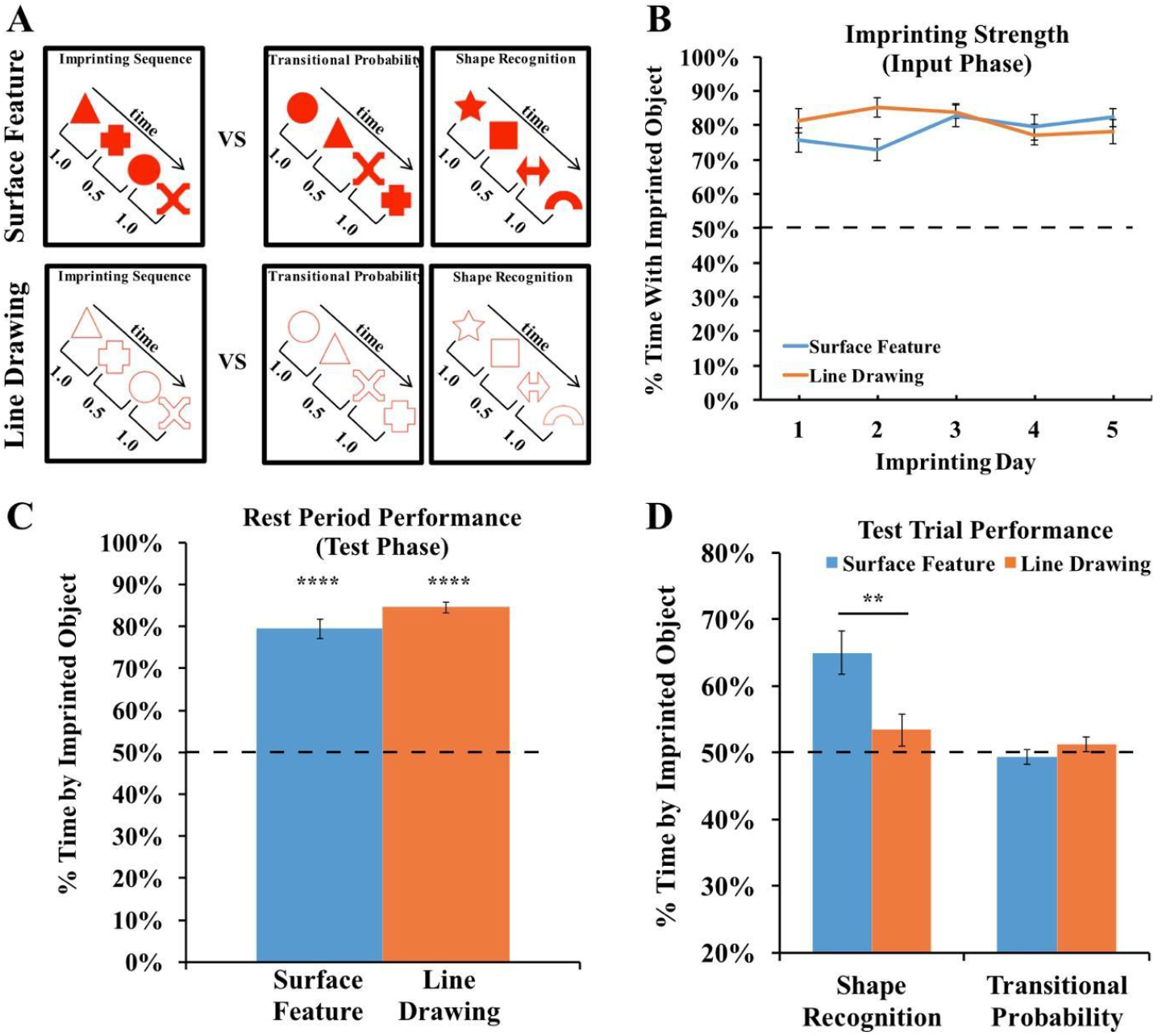
**(A)** Experiment 3 method. During the input phase, an imprinting sequence defined by the transitional probabilities (TPs) within and between shape pairs appeared on one display wall at a time. The imprinting sequence either contained shapes with surface features or line drawings of shapes. During the test phase, we presented chicks with two-alternative forced-choice tasks. In each test trial, one display wall showed the imprinting sequence, and the other display wall showed either the same shapes as in the imprinting sequence but in novel orders (TP trials) or a sequence of novel shapes (Shape Recognition trials). **(B)** Strength of the imprinting response in Experiment 3 during the input phase. **(C)** Strength of the imprinting response in Experiment 3 during the rest periods of the test phase. The chicks imprinted equally strongly across the Surface Feature and Line Drawing Conditions. **(D)** Recognition performance in the Shape Recognition and Transitional Probability Conditions. The chicks showed superior object recognition performance in the Surface Feature Condition compared to the Line Drawing Condition.

For the Surface Feature Condition, we used the data previously reported in Wood, Johnson, & Wood (2019). In that paper, we tested whether newborn chicks can encode the transitional probabilities (TPs) between shapes in a sequence. During the input phase, the chicks were reared with an imprinting sequence consisting of a stream of four shapes, and the order of the shapes was defined by the TPs within and between shape pairs. During the test phase, we presented two types of test trials. On the shape recognition trials, one monitor showed a sequence of familiar shapes, and the opposite monitor showed a sequence of novel shapes. On the TP trials, both monitors showed the familiar shapes, but we manipulated the TPs between shapes. One monitor showed a familiar TP sequence, in which the TPs between shapes matched the imprinting sequence, and the opposite monitor showed a novel TP sequence, in which the TPs between shapes did not match the imprinting sequence. In the original study, we found that the chicks successfully distinguished between the sequences on the shape recognition trials, but failed to distinguish between the sequences on the TP trials. Here, we repeated this experiment with one crucial change: rather than presenting sequences of shapes with surface features, we presented sequences of red line drawing shapes (Figure 6A).

### A. Method

For a detailed description of the methods, see Wood et al. (2019). In the present study, we used a similar design as the original study, except that the chicks were imprinted and tested with red line drawings, rather than red objects with surface features. As in the original study, we tested the chicks with both shape recognition and TP test trials. We tested 12 subjects in the Surface Feature Condition (Wood et al. 2019) and 10 subjects in the Line Drawing Condition. No subjects were excluded from the analyses.

### B. Results and Discussion

We first examined the strength of the imprinting response to ensure that the chicks imprinted equally strongly across the two conditions. As shown in Figure 6B, the chicks in both conditions spent the majority of their time in proximity to the imprinted object during the input phase (one-sample *t*-tests, Surface Feature Condition: *t*(11) = 12.35, *p* < 10^−7^, Cohen’s *d* = 3.57; Line Drawing Condition: *t*(11) = 18.77, *p* < 10^−8^, Cohen’s *d* = 5.42). Unlike in Experiments 1 and 2, the chicks did not show a stronger imprinting response in the Surface Feature Condition than the Line Drawing Condition (independent samples *t*-test, *t*(22) = 0.7, *p* = .94, Cohen’s *d* = 0.03). Similarly, as shown in Figure 6C, the chicks in both conditions spent the majority of their time in proximity to the imprinted object during the rest periods (one-sample *t*-tests, Surface Feature Condition: *t*(11) = 13.06, *p* < 10^−7^, Cohen’s *d* = 3.77; Line Drawing Condition: *t*(9) = 28.59, *p* < 10^−10^, Cohen’s *d* = 9.04). Unlike in Experiments 1 and 2, the chicks did not show a stronger imprinting response during the rest periods in the Surface Feature Condition than the Line Drawing Condition (independent samples *t*-test, *t*(20) =1.69, *p* = .11, Cohen’s *d* = 0.75). Together, these analyses show that the chicks did not imprint more strongly in the Surface Feature Condition, allowing for a more direct comparison of recognition performance across the conditions.

The chicks’ recognition performance is shown in Figure 6D. An ANOVA with the within-subjects factor of Trial Type (Shape Recognition vs. TP trials) and the between-subjects factor of Condition (Surface Feature vs. Line Drawing) revealed a significant main effect of Trial Type (*F*(1,20) = 15.95, *p* = .001, η_p_ ^2^ = .44), a significant main effect of Condition (*F*(1,20) = 4.38, *p* = .049, η_p_ ^2^ = .18), and a significant interaction between Trial Type and Condition (F(1,20) = 9.16, p=.007,ηp2 = .31).

When reared with a sequence of shapes containing surface features, the chicks could reliably distinguish between familiar shapes and novel shapes (one-sample *t*-test, *t*(11) = 4.67, *p* = .0007, Cohen’s *d* = 1.35). In contrast, when reared with a sequence of line drawing shapes, the chicks failed to distinguish between familiar shapes and novel shapes (one-sample *t*-test, *t*(9) = 1.30, *p* = .22, Cohen’s *d* = 0.41). As in the original study (Wood et al., 2019), the chicks in both conditions failed to distinguish between the sequences based on the TPs between shapes (Surface Feature Condition: *t*(11) = 0.56, *p* = 0.59, Cohen’s *d* = -0.16; Line Drawing Condition: *t*(9) = 1.04, *p* = .32, Cohen’s *d* = 0.33).

On the individual-subject level, 8 of the 12 chicks in the Surface Feature Condition showed a statistically significant preference for the familiar shapes (7 chicks: *p* < .0001; 1 chick: *p* < .05). In contrast, in the Line Drawing Condition, only one chick exceeded chance level on the shape recognition trials, while one other chick performed significantly below chance level. As in Experiments 1 and 2, newborn chicks developed superior object recognition performance when reared with objects containing surface features versus line drawings.

## V. GENERAL DISCUSSION

A deep understanding of object recognition requires understanding the role of visual experience in development. Here, we reveal a set of conditions under which object recognition fails to emerge in newborn animals: when a newborn’s visual experience with objects consists solely of line drawings. When newborn chicks were reared with objects containing surface features, the chicks developed robust view-invariant object recognition. In contrast, when chicks were reared with line drawings of objects, the chicks failed to develop object recognition. Notably, the chicks reared with the line drawings performed at chance level, despite acquiring over 100 hours of experience with the objects. Thus, the development of object recognition requires visual experience with the surface features of objects.

Interestingly, the chicks reared with line drawings failed to build accurate object representations despite being raised in environments that contained some surface features. The walls and floor of the chamber contained surface features, as did the heaps of grain consumed during feeding. Nevertheless, when the *objects* in the chicks’ visual environment lacked surface features, the chicks failed to build accurate representations. This finding suggests that experience with surface features per se is not sufficient for the development of object recognition; rather, newborn visual systems need experience with the surface features of objects.

These results add to a growing body of work mapping out the conditions under which object recognition does, and does not, emerge in newborn animals. For instance, studies with newborn chicks have revealed two constraints on the development of object recognition. First, there is a “slowness constraint” on newborn vision: object recognition emerges when newborn chicks are reared with slowly moving objects, but not quickly moving objects (Wood & Wood, 2016b). When chicks are reared with quickly moving objects, their object representations become distorted in the direction of object motion and fail to generalize to novel viewpoints and rotation speeds. Second, there is a “smoothness constraint” on newborn vision: object recognition emerges when newborn chicks are reared with temporally smooth objects, but not temporally non-smooth objects (Wood, 2016; Wood et al., 2016). When chicks are reared with temporally non-smooth objects, their object representations are less selective for object identity. The present study extends this literature by demonstrating that experience with slow and smooth objects is not sufficient for the development of object recognition. The line drawings in Experiments 1-3 moved slowly and smoothly over time, but the chicks nevertheless failed to develop object recognition. Together, these findings indicate that the development of object recognition requires experience with naturalistic objects: objects that have surface features and move slowly and smoothly over time.

More generally, these results build on a large body of research using animal models to examine the mechanisms of object recognition and early visual learning. For decades, newborn chicks have been used to characterize the effects of visual experience on the brain (e.g., Bateson, Horn, & Rose, 1972; Horn, McCabe, & Bateson, 1979; Horn, 1981) and to isolate the neural mechanisms that underlie imprinting (e.g., McCabe, Horn, & Bateson, 1981; McCabe, Cipolia-Neto, Horn, & Bateson, 1982). Studies of chicks have also revealed predispositions that might shape early visual learning (e.g., Lamaire, 2020; Versace, Martinho-Truswell, Kacelnik, & Vallortigara, 2018; Wood, 2017). Another important animal model for studying early visual learning is rodents. Studies of rats provide converging evidence that high-level vision is not unique to primates. Like newborn chicks, rats can recognize objects across novel viewpoints (e.g., Zoccolan, Oertelt, DiCarlo, & Cox, 2009). Normal visual development in rats also requires experience with a slow and smooth visual world (Matteucci & Zoccolan, 2020), suggesting that avian and mammalian brains are subject to common developmental constraints. Specifically, when newborn rats were reared with frame-scrambled versions of natural movies (which preserved the natural spatial statistics but resulted in quickly changing, temporally unstructured input), the rats developed fewer complex cells in primary visual cortex, the cells showed abnormally fast response dynamics, and the cells were less likely to support stable decoding of stimulus orientation. Thus, depriving newborn animals of slowly changing visual experiences disrupts normal visual development in both birds and mammals, potentially reflecting a shared cortex-like canonical circuit found across taxa (Stacho et al., 2020).

### A. Limitations of this study and directions for future research

While these results contribute to our understanding of early visual development, there are limitations to this work that will require additional future research. First, these chicks were reared with either one 3D object (Experiments 1 and 2) or four 2D objects (Experiment 3). It is therefore possible that chicks could develop object recognition in a visual world consisting solely of line drawings if there were more line drawings in the environment and/or if those line drawings were more complex (e.g., line drawings containing polyhedral shapes that included L, Y, and T junctures between lines). Future studies could distinguish between these possibilities by rearing chicks in more complex “line drawing worlds.”

Second, we used the chicks’ preference for their imprinted object as a measure of object recognition performance. While successful performance provides evidence for object recognition (e.g., in the surface feature conditions), the absence of a preference (e.g., in the line drawing conditions) does not necessarily provide evidence for a lack of object recognition. For instance, it is possible that the chicks in the line drawing conditions perceived the test objects as different but grouped them in the same object category. Of course, this alternative explanation must then explain why the presence of surface features would lead chicks to categorize objects differently from one another, whereas line drawings would lead chicks to categorize objects together. It would be interesting for future studies to test chicks using alternative methods (e.g., reinforcement learning) to explore whether the present findings generalize to other object recognition tasks.

Third, these results do not reveal *why* surface features are necessary for the development of object recognition. Why do newborn chicks fail to understand line drawings, when mature animals (including birds) can readily recognize objects presented in line drawings? One possibility is that the mechanisms underlying object recognition require patterned input from natural visual objects in order to develop a receptive field structure that efficiently recovers edges and lines. Specifically, in natural visual environments, the edges of objects and surfaces are typically marked by discrete changes in surface attributes, and mature visual systems contain neurons tuned to the orientation of these contours, responding to edges and lines (Hubel & Wiesel, 1962; 1968). To develop these orientation-tuned detectors, newborn brains may require visual experience of objects with surface features.

Moreover, line drawings are impoverished compared to real objects, and the surface features that appear on real objects may provide valuable information for building accurate representations of an object’s three-dimensional shape. For example, the surface features on the objects used in Experiments 1 and 2 had gradients of luminance that moved as the object rotated, creating flow field cues for three-dimensional shape. Accordingly, it is possibility that experience with realistic objects is necessary to develop the ability to recognize objects in line drawings. Future controlled-rearing experiments could test this hypothesis directly by examining whether experience with realistic objects allows for the development of line drawing understanding.

Ultimately, a deep mechanistic understanding of the role of experience in the development of object recognition will require task-performing computational models that can simulate the complex interactions between newborn brains and the visual environment. The present results should be valuable for this enterprise because they provide precise descriptions of how specific visual inputs relate to specific object recognition outputs in a newborn model system. These input-output patterns can serve as benchmarks for measuring the accuracy of computational models. Specifically, to explain the development of object recognition, a computational model would need to produce two patterns. First, the model should successfully develop view-invariant object recognition when trained with realistic objects that move slowly and smoothly over time. Second, the model should fail to develop object recognition when trained solely with line drawings.

### B. Conclusion

The present study provides evidence that the development of object recognition requires experience with the surface features of objects. Newborn chicks develop enhanced object recognition performance when reared with objects containing surface features compared to line drawings. This study sheds light on how a fundamental ability emerges in newborn animals and provides precise input-output patterns for measuring the accuracy of task-performing computational models of visual development.

## Acknowledgements

This research was funded by NSF CAREER Grant BCS-1351892 and a James S. McDonnell Foundation Understanding Human Cognition Scholar Award. This research was conducted at the University of Southern California.

## Footnotes

1 Human infants’ enhanced attention to lines that depict corners and edges might reflect either an understanding of line drawings or be driven by more basic perceptual preferences underlying infant vision (Hayden, Bhatt, & Quinn, 2006).

2 The data from the baseline (surface feature) conditions in Experiments 1, 2, & 3 were published previously in Wood (2013), Wood & Wood (2016), and Wood, Johnson, & Wood (2019), respectively. In the present study, we directly contrasted chicks reared with line drawings of objects versus objects with surface features.

## References

Bateson, P. P. G., & Jaeckel, J. B. (1976). Chicks’ preferences for familiar and novel conspicuous objects after different periods of exposure. Animal Behaviour, 24, 386–390.

Bateson, P. P., Horn, G., & Rose, S. P. (1972). Effects of early experience on regional incorporation of precursors into RNA and protein in the chick brain. Brain Research, 39(2), 449–465.

Biederman, I. (1987). Recognition-by-components: a theory of human image understanding. Psychological Review, 94(2), 115–147.

Biederman, I., & Ju, G. (1988). Surface versus edge-based determinants of visual recognition. Cognitive Psychology, 20(1), 38–64.

Clottes, J. (2000). Heilbrunn Timeline of Art History. Chauvet Cave (ca. 30,000 BC).

DiCarlo, J. J., Zoccolan, D., & Rust, N. C. (2012). How does the brain solve visual object recognition? Neuron, 73(3), 415–434.

Goodnow, J. J. (1977). Children’s Drawing. Cambridge, MA. Harvard University Press.

Güntürkün, O., & Bugnyar, T. (2016). Cognition without cortex. Trends in Cognitive Sciences, 20(4), 291–303.

Ham, A.D. & Osorio, D. (2007). Colour preferences and colour vision in poultry chicks. Proceedings of the Royal Society: B., 274, 1941–1948.

Hayward, W. G., & Williams, P. (2000). Viewpoint dependence and object discriminability. Psychological Science, 11(1), 7–12.

Horn, G. (1981). Review Lecture: Neural mechanisms of learning: an analysis of imprinting in the domestic chick. Proceedings of the Royal Society: B. 213, 101–137.

Horn, G. (2004). Pathways of the past: the imprint of memory. Nature Reviews Neuroscience, 5(2), 108–120.

Horn, G., McCabe, B., & Bateson, P. (1979). Autoradiographic study of the chick brain after imprinting. Brain Research, 168, 361–373.

Hubel, D. H., & Wiesel, T. N. (1962). Receptive fields, binocular interaction and functional architecture in cats visual cortex. The Journal of Physiology, 160(1), 106–154.

Hubel, D. H., & Wiesel, T. N. (1968). Receptive fields and functional architecture of monkey striate cortex. The Journal of Physiology, 195(1), 215–243.

Ishai, A., Ungerleider, L. G., Martin, A., & Haxby, J. V. (2000). The representation of objects in the human occipital and temporal cortex. Journal of Cognitive Neuroscience, 12, 35–51.

Itakura, S. (1994). Recognition of line-drawing representations by a chimpanzee (Pan troglodytes). The Journal of General Psychology, 121(3), 189–197.

Jarvis, E. D., Güntürkün, O., Bruce, L., Csillag, A., Karten, H., Kuenzel, W., et al. (2005). Avian brains and a new understanding of vertebrate brain evolution. Nature Reviews Neuroscience, 6(2), 151–159.

Johnson, M., Bolhuis, J., & Horn, G. (1985). Interaction between acquired preferences and developing predispositions during imprinting. Animal Behaviour, 33, 1000–1006.

Karten, H. J. (2013). Neocortical evolution: neuronal circuits arise independently of lamination. Current Biology, 23(1), R12–5.

Kennedy, J. M., & Ross, A. S. (1975). Outline picture perception by the Songe of Papua. Perception, 4(4), 391–406.

Lemaire, B.S. (2020). No evidence of spontaneous preference for slowly moving objects in visually naïve chicks. Scientific Reports, 10, 6277.

Matteucci, G., & Zoccolan, D. (2020). Unsupervised experience with temporal continuity of the visual environment is causally involved in the development of V1 complex cells. Science Advances, 6(22), eaba3742. DOI: 10.1126/sciadv.aba3742

McCabe, B. J., Horn, G., & Bateson, P. P. G. (1981). Effects of restricted lesions of the chick forebrain on the acquisition of filial preferences during imprinting. Brain Research, 205, 29–37.

McCabe, B. J., Cipolla-Neto, J., Horn, G., & Bateson, P. (1982). Amnesic effects of bilateral lesions placed in the hyperstriatum ventrale of the chick after imprinting. Experimental Brain Research, 48, 13–21.

Miura, M., Nishi, D. & Matsushima, T. (2020). Combined predisposed preferences for colour and biological motion make robust development of social attachment through imprinting. Animal Cognition, 23, 169–188.

Naor-Raz, G., Tarr, M. J., & Kersten, D. (2003). Is color an intrinsic property of object representation? Perception, 32(6), 667–680.

Nakamori, T., Maekawa, F., Sato, K., Tanaka, K., & Ohki-Hamazaki, H. (2013). Neural basis of imprinting behavior in chicks. Dev. Growth Differ. 55, 198–206. doi: 10.1111/dgd.12028

Prasad, A., Wood, S. M. W., & Wood, J. N. (2019). Using automated controlled rearing to explore the origins of object permanence. Developmental Science, 22(3), e12796.

Price, C. J., & Humphreys, G. W. (1989). The effects of surface detail on object categorization and naming. The Quarterly Journal of Experimental Psychology Section A, 41(4), 797–828.

Rossion, B., & Pourtois, G. (2004). Revisiting Snodgrass and Vanderwart’s object pictorial set: The role of surface detail in basic-level object recognition. Perception, 33(2), 217–236.

Stacho, M. et al. (2020). A cortex-like canonical circuit in the avian forebrain. Science, 369(6511), eabc5534.

Sayim, B., & Cavanagh, P. (2011). What line drawings reveal about the visual brain. Frontiers in Human Neuroscience, 5.

Spelke, E. (1990). Principles of object perception. Cognitive Science, 14(1), 29–56.

Tanaka, M. (2006). Recognition of pictorial representations by chimpanzees (Pan troglodytes). Animal Cognition, 10(2), 169–179.

Tarr, M. J., Kersten, D., & Bulthoff, H. H. (1998). Why the visual recognition system might encode the effects of illumination. Vision Research, 38(15-16), 2259–2275.

Versace, E., Martinho-Truswell, A., Kacelnik, A., & Vallortigara, G. (2018). Priors in animal and artificial intelligence: where does learning begin? Trends in Cognitive Sciences, 22(11), 963–965

Vuong, Q. C., Peissig, J. J., Harrison, M. C., & Tarr, M. J. (2005). The role of surface pigmentation for recognition revealed by contrast reversal in faces and Greebles. Vision Research, 45(10), 1213–1223.

Walther, D. B., Chai, B., Caddigan, E., Beck, D. M., & Fei-Fei, L. (2011). Simple line drawings suffice for functional MRI decoding of natural scene categories. Proceedings of the National Academy of Sciences of the United States of America, 108(23), 9661–9666.

Wasserman, E. A., Gagliardi, J. L., Cook, B. R., Kirkpatrick-Steger, K., Astley, S. L., & Biederman, I. (1996). The pigeon’s recognition of drawings of depth-rotated stimuli. Journal of Experimental Psychology: Animal Behavior Processes, 22(2), 205–221.

Wood, J. N. (2013). Newborn chickens generate invariant object representations at the onset of visual object experience. Proceedings of the National Academy of Sciences, 110(34), 14000–14005.

Wood, J. N. (2014). Newly hatched chicks solve the visual binding problem. Psychological Science, 25(7), 1475–1481.

Wood, J. N. (2015). Characterizing the information content of a newly hatched chick’s first visual object representation. Developmental Science, 18(2), 194–205.

Wood, J. N. (2016). A smoothness constraint on the development of object recognition. Cognition, 153, 140–145.

Wood, J. N., Prasad, A., Goldman, J. G., & Wood, S. M. W. (2016). Enhanced learning of natural visual sequences in newborn chicks. Animal Cognition, 19(4), 835–845.

Wood, S. M. W., & Wood, J. N. (2015). A chicken model for studying the emergence of invariant object recognition. Frontiers in Neural Circuits, 9(89), 7.

Wood, J. N., & Wood, S. M. W. (2016). The development of newborn object recognition in fast and slow visual worlds. Proceedings of the Royal Society: Biological Sciences, 283(1829).

Wood, J. N. (2017). Spontaneous preference for slowly moving objects in visually naïve animals. Open Mind, 1(2), 111–122. doi: 10.1162/OPMI_a_00012.

Wood, J. N., & Wood, S. M. W. (2017). Measuring the speed of newborn object recognition in controlled visual worlds. Developmental Science. http://doi.org/10.1111/desc.12470

Wood, J. N., & Wood, S. M. W. (2018). The development of invariant object recognition requires visual experience with temporally smooth objects. Cognitive Science, 39(17), 2885–16.

Wood, S. M. W., & Wood, J. N. (2019). Using automation to combat the replication crisis: A case study from controlled-rearing studies of newborn chicks. Infant Behavior and Development, 57, 101329.

Wurm, L. H., Legge, G. E., Isenberg, L. M., & Luebker, A. (1993). Color improves object recognition in normal and low vision. Journal of Experimental Psychology: Human Perception and Performance, 19(4), 899–911.

Yonas, A., & Arterberry, M. E. (1994). Infants perceive spatial structure specified by line junctions. Perception, 23(12), 1427–1435.

Zoccolan, D., Oertelt, N., DiCarlo, J. J., & Cox, D. D. (2009). A rodent model for the study of invariant visual object recognition. Proceedings of the National Academy of Sciences, 106(21), 8748–8753.

